# An engineered decoy receptor for SARS-CoV-2 broadly binds protein S sequence variants

**DOI:** 10.1101/2020.10.18.344622

**Authors:** Kui K. Chan, Timothy J.C. Tan, Krishna K. Narayanan, Erik Procko

## Abstract

The spike S of SARS-CoV-2 recognizes ACE2 on the host cell membrane to initiate entry. Soluble decoy receptors, in which the ACE2 ectodomain is engineered to block S with high affinity, potently neutralize infection and, due to close similarity with the natural receptor, hold out the promise of being broadly active against virus variants without opportunity for escape. Here, we directly test this hypothesis. We find an engineered decoy receptor, sACE2_2_.v2.4, tightly binds S of SARS-associated viruses from humans and bats, despite the ACE2-binding surface being a region of high diversity. Saturation mutagenesis of the receptor-binding domain (RBD) followed by in vitro selection, with wild type ACE2 and the engineered decoy competing for binding sites, failed to find S mutants that discriminate in favor of the wild type receptor. Variant N501Y in the RBD, which has emerged in a rapidly spreading lineage (B.1.1.7) in England, enhances affinity for wild type ACE2 20-fold but remains tightly bound to engineered sACE22.v2.4. We conclude that resistance to engineered decoys will be rare and that decoys may be active against future outbreaks of SARS-associated betacoronaviruses.

## INTRODUCTION

Zoonotic coronaviruses have crossed over from animal reservoirs multiple times in the past two decades, and it is almost certain that wild animals will continue to be a source of devastating outbreaks. Unlike ubiquitous human coronaviruses responsible for common respiratory illnesses, these zoonotic coronaviruses with pandemic potential cause serious and complex diseases, in part due to their tissue tropisms driven by receptor usage. Severe Acute Respiratory Syndrome Coronaviruses 1 (SARS-CoV-1) and 2 (SARS-CoV-2) engage angiotensin-converting enzyme 2 (ACE2) for cell attachment and entry (*1*-*7*). ACE2 is a protease responsible for regulating blood volume and pressure that is expressed on the surface of cells in the lung, heart and gastrointestinal tract, among other tissues (*8, 9*). The ongoing spread of SARS-CoV-2 and the disease it causes, COVID-19, has had a crippling toll on global healthcare systems and economies, and effective treatments and vaccines are urgently needed.

As SARS-CoV-2 becomes endemic in the human population, it has the potential to mutate and undergo genetic drift and recombination. To what extent this will occur as increasing numbers of people are infected and mount counter immune responses is unknown, but already a variant in the viral spike protein S (D614G) has rapidly emerged from multiple independent events and effects S protein stability and dynamics (*10, 11*). Another S variant (D839Y) became prevalent in Portugal, possibly due to a founder effect (*12*). SARS-CoV-2 has a moderate mutation rate estimated at 10^−3^ substitutions per site per year (*13*). However, the virus has undergone rapid mutation and adaptation after infecting mink in Denmark, from which it then crossed back to humans (*14*), causing Danish authorities to order 17 million farmed mink culled to pre-emptively prevent the possible emergence of vaccine-resistant variants. Additionally, large changes in coronavirus genomes have frequently occurred in nature from recombination events, especially in bats where co-infection levels can be high (*15, 16*). Recombination of MERS-CoVs has been documented in camels (*17*), there are reported cases of recombination between co-circulating SARS-CoV-2 variants (*18*), and SARS-CoV-2 itself may have emerged through recombination of coronavirus genomes (*19*). This will all have profound implications for the current pandemic’s trajectory, the potential for future coronavirus pandemics, and whether drug or vaccine resistance in SARS-CoV-2 emerges and becomes widespread.

The viral spike is a vulnerable target for neutralizing monoclonal antibodies that are progressing through clinical trials, yet in tissue culture escape mutations in the spike rapidly emerge to all antibodies tested (*20*). Deep mutagenesis of the isolated receptor-binding domain (RBD) by yeast surface display has easily identified mutations in S that retain high expression and ACE2 affinity, yet are no longer bound by monoclonal antibodies and confer resistance (*21*). This has motivated the development of cocktails of non-competing monoclonals (*20, 22*), inspired by lessons learned from the treatment of HIV-1 and Ebola, to limit the possibilities for the virus to escape. Notably, drug maker Eli Lilly has a monoclonal monotherapy (LY-CoV555) in advanced trials (NCT04427501) where the selection of resistant virus variants in patients has occurred. A trial update added an arm with a second monoclonal (LY-CoV016) and the company has not reported putative resistance variants in patients receiving the cocktail thus far. However, even the use of monoclonal cocktails does not address future coronavirus spill overs from wild animals that may be antigenically distinct. Indeed, large screening efforts were required to find antibodies from recovered SARS-CoV-1 patients that cross-react with SARS-CoV-2 (*23*), indicating antibodies have confined capacity for interacting with variable epitopes on the spike surface, and are unlikely to be broad and pan-specific for all SARS-related viruses.

An alternative protein-based antiviral to monoclonal antibodies is to use soluble ACE2 (sACE2) as a decoy to compete for receptor-binding sites on the viral spike (*6, 24*-*27*). In principle, the virus has limited potential to escape sACE2-mediated neutralization without simultaneously decreasing affinity for the native ACE2 receptor, rendering the virus less virulent. Wild type sACE2 is currently in a phase II clinical trial (*28*) and multiple groups have now engineered sACE2 to create high affinity decoys for SARS-CoV-2 that rival matured monoclonal antibodies for potent neutralization of infection (*27, 29, 30*). In our group, deep mutagenesis was used to identify a large number of mutations in ACE2 that increase affinity for S (*27*). These mutations were dispersed across the interface and also at distal sites where they are predicted to enhance folding of the virus-recognized conformation. A combination of three mutations, called sACE2_2_.v2.4, increases affinity 35-fold and binds SARS-CoV-2 S (K_D_ 600 pM) with affinity comparable to the best monoclonal antibodies (*27*). Even tighter apparent affinities are reached through avid binding to trimeric spike expressed on a membrane. Despite engineering being focused exclusively on SARS-CoV-2 affinity, sACE2_2_.v2.4 potently neutralized authentic SARS-CoV-1 and −2 infection in tissue culture, suggesting it’s close resemblance to the wild type receptor confers broad activity against ACE2-utilizing betacoronaviruses generally. Soluble ACE2_2_.v2.4 is dimeric and monodisperse without aggregation, catalytically active, highly soluble, stable after storage at 37°C for days, and well expressed at levels greater than the wild type protein. Due to both its high activity and favorable properties for manufacture, sACE2_2_.v2.4 is a genuine drug candidate for preclinical development.

Engineered, high affinity decoy receptors, while very similar to natural ACE2, nonetheless have mutations present at or near the interaction surface. There is therefore an opportunity for viral spike variants to discriminate between an engineered decoy and wild type receptors, providing a route towards resistance. Here, we show that the engineered decoy sACE2_2_.v2.4 binds broadly and tightly to the RBDs of diverse SARS-associated betacoronaviruses that use ACE2 for entry. We further fail to find mutations within the RBD, which directly contacts ACE2 and is where possible escape mutations will most likely reside, that redirect specificity towards the wild type receptor. We conclude that resistance to an engineered decoy receptor will be rare, and sACE2_2_.v2.4 targets common attributes for affinity to S in SARS-associated viruses.

## RESULTS

### An engineered decoy receptor broadly binds RBDs from SARS-associated CoVs with tight affinity

The affinities of the decoy receptor sACE2_2_.v2.4 were determined for purified RBDs from the S proteins of five coronaviruses from *Rhinolophus* bat species (isolates LYRa11, Rs4231, Rs7327, Rs4084 and RsSHC014) and two human coronaviruses, SARS-CoV-1 and SARS-CoV-2. These viruses fall within a common clade of betacoronaviruses that have been experimentally validated to use human ACE2 as an entry receptor (*7*). They share close sequence identity within the RBD core while variation is highest within the functional ACE2 binding site (Figures 1 and S1), possibly due to a co-evolutionary ‘arms race’ with polymorphic ACE2 sequences in ecologically diverse bat species (*31*). Affinity was measured by biolayer interferometry (BLI), with sACE2_2_ (a.a. S19-G732) fused at the C-terminus with the Fc moiety of human IgG1 immobilized to the sensor surface and monomeric 8his-tagged RBD (Figure S2) used as the soluble analyte. This arrangement excludes avidity effects, which otherwise cause artificially tight (picomolar) apparent affinities whenever dimeric sACE2_2_ in solution is bound to immobilized RBD decorating an interaction surface. Wild type sACE2_2_ bound all the RBDs with affinities ranging from 16 nM for SARS-CoV-2 to 91 nM for LYRa11, with median affinity 60 nM (Table 1). The measured affinities for the RBDs of SARS-CoV-1 and SARS-CoV-2 are comparable to published data (*4, 27, 32*-*34*). Engineered sACE2_2_.v2.4 displayed large increases in affinity for all the RBDs, with K_D_s ranging from 0.4 nM for SARS-CoV-2 to 3.5 nM for isolate Rs4231, with median affinity less than 2 nM (Table 1). The approximate 35-fold affinity increase of the engineered decoy applies universally to coronaviruses in the test panel and the molecular basis for affinity enhancement must therefore be grounded in common attributes of RBD/ACE2 recognition.

**Figure 1.**
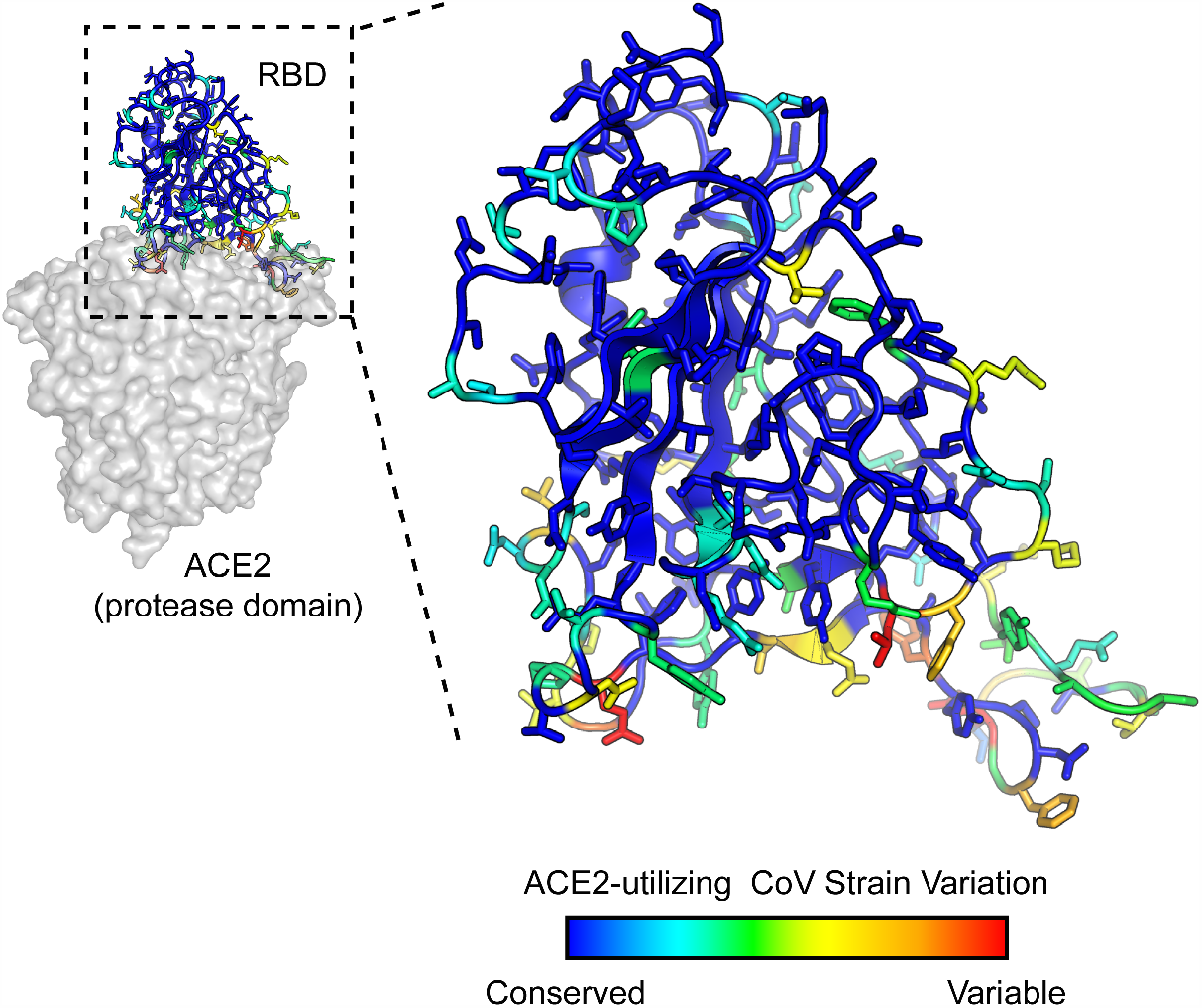
SARS-associated coronaviruses have high sequence diversity at the ACE2-binding site. The RBD of SARS-CoV-2 (PDB 6M17) is colored by diversity between 7 SARS-associated CoV strains (blue, conserved; red, variable).

**Table 1.** BLI kinetics for immobilized sACE2_2_-IgG1 binding to coronavirus RBDs.

|  |  |  |  |  |  |  |  |  |
| --- | --- | --- | --- | --- | --- | --- | --- | --- |
| CoV strain <sup>a</sup> | Wild type sACE2<sub>2</sub>-IgG1 <sup>b</sup> |  |  |  | sACE2<sub>2</sub>.v2.4-IgG1 |  |  |  |
|  | k<sub>on</sub> (M<sup>-1</sup> s<sup>-1</sup>) | k<sub>off</sub> (s<sup>-1</sup>) | K<sub>D</sub> (nM) | χ<sup>2</sup> <sup>c</sup> | k<sub>on</sub> (M<sup>-1</sup> s<sup>-1</sup>) | k<sub>off</sub> (s<sup>-1</sup>) | K<sub>D</sub> (nM) | χ<sup>2</sup> |
| LYRa11 | 8.7 × 10<sup>5</sup> | 7.9 × 10<sup>-2</sup> | 91 | 0.12 | 1.4 × 10<sup>6</sup> | 2.5 × 10<sup>-3</sup> | 1.8 | 0.10 |
| Rs7327 | 6.4 × 10<sup>5</sup> | 4.0 × 10<sup>-2</sup> | 63 | 0.25 | 9.8 × 10<sup>5</sup> | 1.8 × 10<sup>-3</sup> | 1.9 | 0.11 |
| Rs4231 | 3.2 × 10<sup>5</sup> | 2.2 × 10<sup>-2</sup> | 69 | 0.04 | 4.5 × 10<sup>5</sup> | 1.6 × 10<sup>-3</sup> | 3.5 | 0.10 |
| Rs4084 | 2.9 × 10<sup>5</sup> | 2.5 × 10<sup>-2</sup> | 85 | 0.24 | 4.8 × 10<sup>5</sup> | 1.5 × 10<sup>-3</sup> | 3.1 | 0.10 |
| RsSHC014 | 8.8 × 10<sup>5</sup> | 2.6 × 10<sup>-2</sup> | 29 | 0.20 | 1.6 × 10<sup>6</sup> | 2.0 × 10<sup>-3</sup> | 1.3 | 0.29 |
| SARS-1 | 6.6 × 10<sup>3</sup> | 1.2 × 10<sup>-4</sup> | 58 | 0.03 | 3.0 × 10<sup>3</sup> | 5.6 × 10<sup>-6</sup> | 2.1 | 0.03 |
| SARS-2 | 1.4 × 10<sup>6</sup> | 8.1 × 10<sup>-3</sup> | 16 | 0.25 | 6.6 × 10<sup>5</sup> | 2.8 × 10<sup>-4</sup> | 0.4 | 0.09 |
| SARS-2 (Y449K) | 2.0 × 10<sup>6</sup> | 9.0 × 10<sup>-2</sup> | 46 | 0.67 | 4.3 × 10<sup>6</sup> | 4.0 × 10<sup>-3</sup> | 0.9 | 0.71 |
| SARS-2 (N501W) | 2.4 × 10<sup>6</sup> | 5.4 × 10<sup>-3</sup> | 2.3 | 0.43 | 3.3 × 10<sup>6</sup> | 2.8 × 10<sup>-4</sup> | 0.1 | 0.23 |
| SARS-2 (N501Y) | 2.2 × 10<sup>6</sup> | 1.8 × 10<sup>-3</sup> | 0.8 | 0.15 | 3.6 × 10<sup>5</sup> | 1.1 × 10<sup>-4</sup> | < 0.1 | 0.24 |
| <sup>a</sup> Purified RBDs at 5 to 7 concentrations were used as the soluble analytes. |  |  |  |  |  |  |  |  |
| <sup>b</sup> IgG1-Fc fused sACE2<sub>2</sub> was immobilized to anti-human IgG Fc capture biosensors. |  |  |  |  |  |  |  |  |
| <sup>c</sup> χ<sup>2</sup> values represent the goodness of curve fitting. Acceptable values were considered to be less than 3. |  |  |  |  |  |  |  |  |

### A deep mutational scan of the RBD in the context of full-length S reveals residues in the ACE2 binding site are mutationally tolerant

To explore potential sequence diversity in S of SARS-CoV-2 that may act as a ‘reservoir’ for drug resistance, the mutational tolerance of the RBD was evaluated by deep mutagenesis (*35*). Saturation mutagenesis was focused to the RBD (a.a. C336-L517) of full-length S tagged at the extracellular N-terminus with a c-myc epitope for detection of surface expression. The spike library, encompassing 3,640 single amino acid substitutions, was transfected in human Expi293F cells under conditions where cells typically acquire no more than a single sequence variant (*36, 37*). The culture was incubated with wild type, 8his-tagged, dimeric sACE2_2_ at a sub-saturating concentration (2.5 nM). Bound sACE2_2_-8h and surface-expressed S were stained with fluorescent antibodies for flow cytometry analysis (Figure 2A). Compared to cells expressing wild type S, the library was poorly expressed, indicating many mutations are deleterious for folding and expression. A cell population was clearly discernable expressing S variants that bind ACE2 with decreased affinity (Figure 2B). After gating for c-myc-positive cells expressing S, cells with high and low levels of bound sACE2_2_ were collected by fluorescence-activated cell sorting (FACS), called the ACE2-High and ACE2-Low populations, respectively (Figure 2C). Both the expression and sACE2_2_ binding signals decreased over minutes to hours during sorting, possibly due to shedding of the S1 subunit. Cells were therefore collected and pooled from three separate FACS experiments for a combined 8 hours sort time.

**Figure 2.**
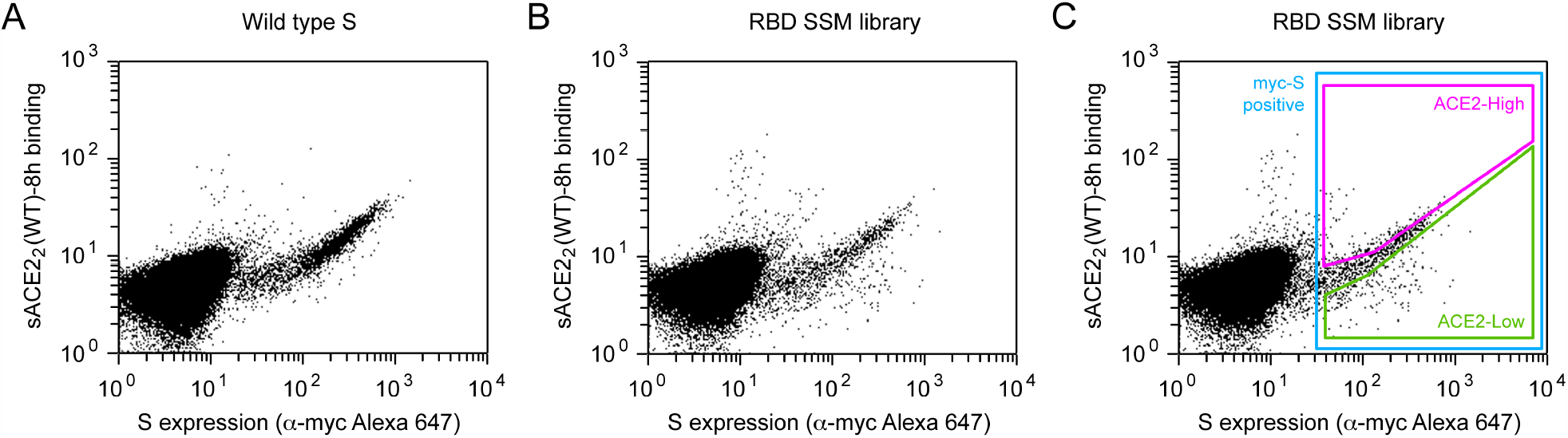
FACS selection for variants of S with high or low binding signal to ACE2. **(A)** Flow cytometry analysis of Expi293F cells expressing full-length S of SARS-CoV-2 with an N-terminal c-myc tag. Staining for the myc-epitope is on the x-axis while the detection of bound sACE2_2_-8h (2.5 nM) is on the y-axis. S plasmid was diluted 1500-fold by weight with carrier DNA so that cells typically express no more than one coding variant; under these conditions most cells are negative. **(B)** Flow cytometry of cells transfected with the RBD single site-saturation mutagenesis (SSM) library shows cells expressing S variants with reduced sACE2_2_-8h binding. **(C)** Gating strategy for FACS. S-expressing cells positive for the c-myc epitope were gated (blue) and the highest (“ACE2-High”) and lowest (“ACE2-Low”) 20% of cells with bound sACE2_2_-8h relative to myc-S expression were collected.

Transcripts in the sorted cells were Illumina sequenced and compared to the naive plasmid library to determine an enrichment ratio for each amino acid substitution (*38*). Mutations in S that express and bind ACE2 tightly are selectively enriched in the ACE2-High sort (Figure S3); mutations that express but have reduced ACE2 binding are selectively enriched in the ACE2-Low sort; and mutations that are poorly expressed are depleted from both sorted populations. Positional conservation scores were calculated by averaging the log_2_ enrichment ratios for each of the possible amino acids at a residue position. By adding conservation scores for both the ACE2-High and ACE2-Low sorts we derive a score for surface expression, which shows that the hydrophobic RBD core is tightly conserved for folding and trafficking of the viral spike (Figure 3A). By comparison, residues on the exposed RBD surface are mutationally permissive for S surface expression. This matches the mutational tolerance of proteins generally.

**Figure 3.**
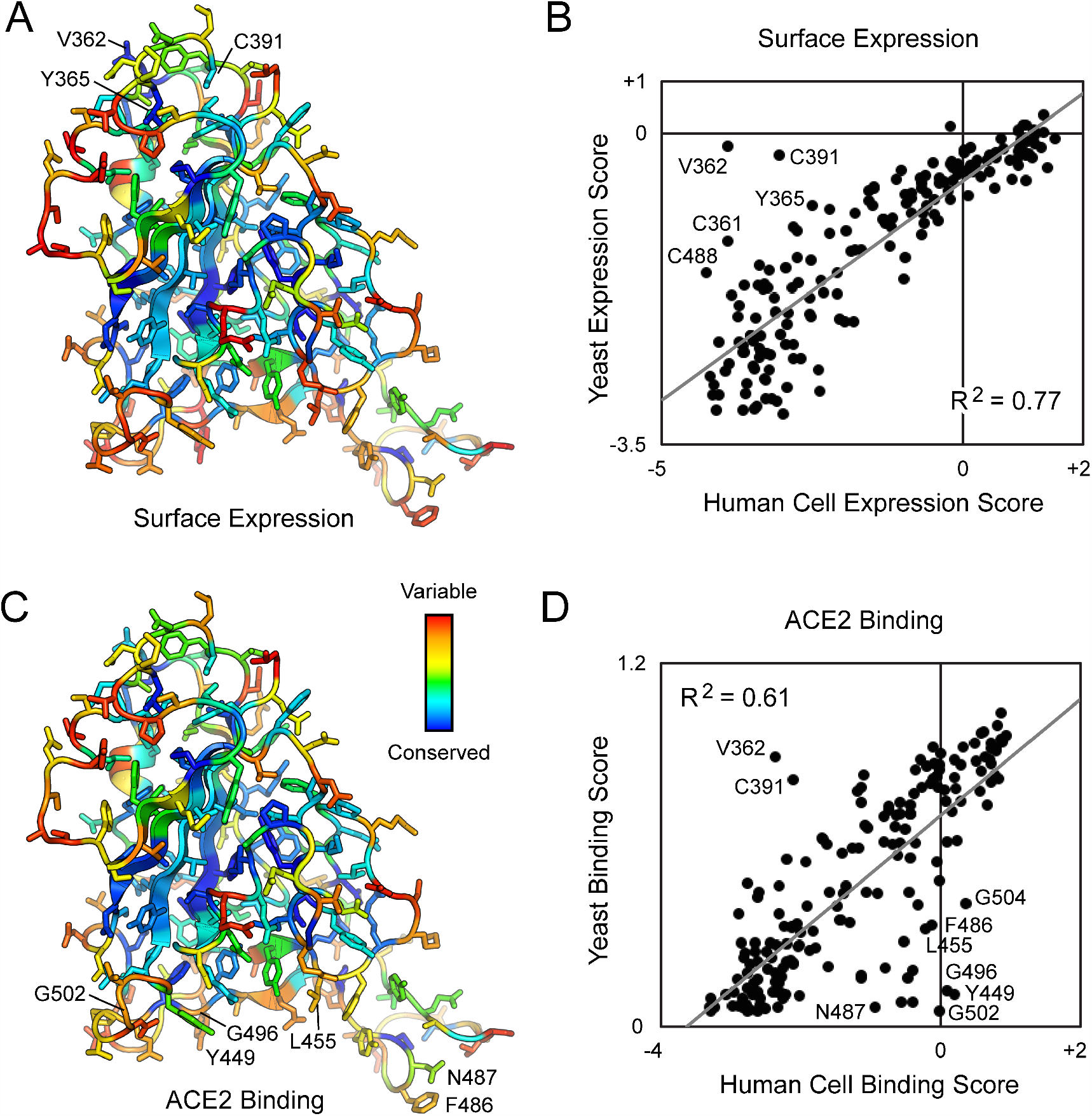
Deep mutagenesis reveals that the ACE2-binding site of SARS-CoV-2 tolerates many mutations. **(A)** Positional scores for surface expression are mapped to the structure of the SARS-CoV-2 RBD (PDB 6M17, oriented as in Figure 1). Blue residues in the protein core are highly conserved in the FACS selection for surface S expression (judged by depletion of mutations from the ACE2-High and ACE2-Low gates), while surface residues in red tolerate mutations. **(B)** Correlation plot of expression scores from mutant selection in human cells of full-length S (x-axis) versus the conservation scores (mean of the log_2_ enrichment ratios at a residue position) from mutant selection in the isolated RBD by yeast display (y-axis). Notable outliers are indicated. **(C)** Conservation scores from the ACE2-High gated cell population are mapped to the RBD structure, with residues colored from low (blue) to high (red) mutational tolerance. **(D)** Correlation plot of RBD conservation scores for high ACE2 binding from deep mutagenesis of S in human cells (x-axis) versus deep mutagenesis of the RBD on the yeast surface (mean of ΔK_D app_; y-axis).

For tight ACE2 binding (i.e. S variants in the ACE2-High population), conservation increases for RBD residues at the ACE2 interface, yet mutational tolerance remains high (Figure 3C). The sequence diversity observed among natural betacoronaviruses, which display high diversity at the ACE2 binding site, is therefore replicated in the deep mutational scan, which predicts the SARS-CoV-2 spike tolerates substantial genetic diversity at the receptor-binding site for function. From this accessible sequence diversity SARS-CoV-2 might feasibly mutate to acquire resistance to monoclonal antibodies or engineered decoy receptors targeting the ACE2-binding site.

There are two ‘hot spot’ regions for interactions at the interface that determine receptor affinity and species adaptation, centered around ACE2 residues K31 and K353 (*39*). These regions are also the locations for substitutions to ACE2 residues T27, L79, and N330 in the engineered sACE2_2_.v2.4 decoy. Mutations to RBD residues at these sites tend to be weakly depleted for high ACE2 binding (Figure S4), but are much more tolerant of mutations than structural positions buried in the RBD core. To highlight a few residues, S-Y505 that packs against the hydrocarbon chain of ACE2-K353 is notably more conserved than most other interfacial residues, with the exceptions of partially buried residues like S-R403 and S-Y453 that likely have additional structural roles; S-Y489 contacting ACE2-T27 and ACE2-K31 has an overall weak preference for aromatic amino acids; and S-G485 on a loop packed against ACE2-L79 has highest tolerance for polar substitutions, possibly to maintain the loop conformation and solubility (Figure S4). Some mutations are found to be highly enriched for ACE2 binding, including small hydrophobic amino acids for S-Q493 and aromatic amino acids for S-N501 that are both anticipated to increase local hydrophobic or aromatic ring packing. This is consistent with observations from yeast surface display of RBD mutants (*40*). That these mutations predicted to increase ACE2 binding are not enriched in circulating SARS-CoV-2 variants suggests the affinity of the virus for its receptor is already sufficient for high transmission and peak fitness (*40*).

### Comparison to a deep mutational scan of the isolated RBD by yeast surface display

Two deep mutational scans have been reported for the isolated RBD displayed on the surface of yeast (*40, 41*). We compare our data, from a selection of full-length S expressed in human cells, to the publicly accessible Starr et al data set (*40*). Important residues within the RBD for surface expression of full-length spike in human cells are closely correlated with data from yeast surface display of the isolated RBD (Figure 3B), with the exception of a notable region. The surface of the RBD opposing the ACE2-binding site (e.g. V362, Y365, and C391) is free to mutate for yeast surface display, but its sequence is constrained in our experiments; this region of the RBD is buried by connecting structural elements to the global fold of an S subunit in the closed-down conformation (this is the dominant conformation for S subunits and is inaccessible to receptor binding; Figure S5) (*2, 4, 42, 43*). While some mutations might have allosteric effects, we note that among substitutions of V362, Y365, and C391, mutations tend to be lightly depleted from the ACE2-Low and heavily depleted from the ACE2-High sorted populations. No mutations at these positions were selectively enriched in just the ACE2-High sort, as might be expected if a mutation favored the open-up conformation through allosteric mechanisms. This is consistent with mutations at these residues reducing ACE2 interactions through defects in folding and decreased surface expression. Compared to single mutations that destabilize the RBD in the closed-down conformation, more extensive engineering with multiple stabilizing mutations has been shown to shift the conformational equilibrium of S subunits to the open-up state (*44*).

We used targeted mutagenesis to individually test alanine substitutions to all the cysteines in the RBD (Figure S5). We found all cysteine-to-alanine mutations severely diminish S surface expression in Expi293F cells, including C391A and C525A on the RBD ‘backside’ that were neutral in the yeast display scan (*40*). These differences demonstrate that there are tighter sequence constraints on the RBD in the context of a full spike expressed at a human cell membrane, yet overall we consider the yeast display and the human cell data sets to closely agree.

For binding to dimeric sACE2_2_, we note that interface residues were more tightly conserved in the Starr et al data set (Figure 3D), possibly a consequence of three differences between the deep mutagenesis experiments. First, our selections for ACE2 binding of S variants at the plasma membrane appears to primarily reflect mutational effects on surface expression, which is almost certainly more stringent in human cells. Yeast permit many poorly folded proteins to leak to the cell surface (*45*). Second, the yeast selections were conducted at multiple sACE2 concentrations from which apparent K_D_ changes were computed (*40*); the Starr et al data in this regard is very comprehensive. Due to the long sort times required for our human cell libraries where only a small fraction of cells express spike, we sorted at a single sACE2_2_ concentration that cannot accurately capture a range of different binding affinities quantitatively. Third, dimeric sACE2_2_ may geometrically complement trimeric S densely packed on a human cell membrane, such that avidity masks the effects of affinity-reducing mutations. Nonetheless, there is overall agreement that ACE2 binding often persists following mutations to the RBD surface, and our data simply suggests mutational tolerance may be even greater than that already observed by Starr et al.

### A screen for S variants that preferentially bind wild type ACE2 over the engineered decoy

Having shown that the ACE2-binding site of SARS-CoV-2 protein S tolerates many mutations, we asked whether mutations might therefore be found that confer resistance to the engineered decoy sACE2_2_.v2.4. Resistance mutations are anticipated to lose affinity for sACE2_2_.v2.4 while maintaining binding to the wild type receptor, and are most likely to reside in the RBD where physical contacts are made. Similar reasoning formed the foundation of a deep mutagenesis-based selection of the isolated RBD by yeast surface display to find escape mutations to monoclonal antibodies, and the results were predictive of escape mutations in pseudovirus growth selections (*21*).

To address whether escape mutations from the engineered decoy might be found in the RBD, we repurposed the S protein library for a specificity selection. Cells expressing the library, encoding all possible substitutions in the RBD, were co-incubated with wild type sACE2_2_ fused to the Fc region of IgG1 and 8his-tagged sACE2_2_.v2.4 at concentrations where both proteins bind competitively (*27*). It was immediately apparent from flow cytometry of the Expi293F culture expressing the S library that there were cells expressing S variants shifted towards preferential binding to sACE2_2_.v2.4, but no significant population with preferential binding to the wild type receptor (Figures 4A and4B). Cells expressing S variants that might preferentially bind sACE2_2_(WT)-IgG1 or sACE2_2_.v2.4 were gated and collected by FACS (Figure 4C), followed by deep sequencing of S transcripts to determine enrichment ratios. There was close agreement between two independent replicate experiments (Figures 4D-4G). Most RBD mutations were depleted following sorting, consistent with deleterious effects on S folding and expression.

**Figure 4.**
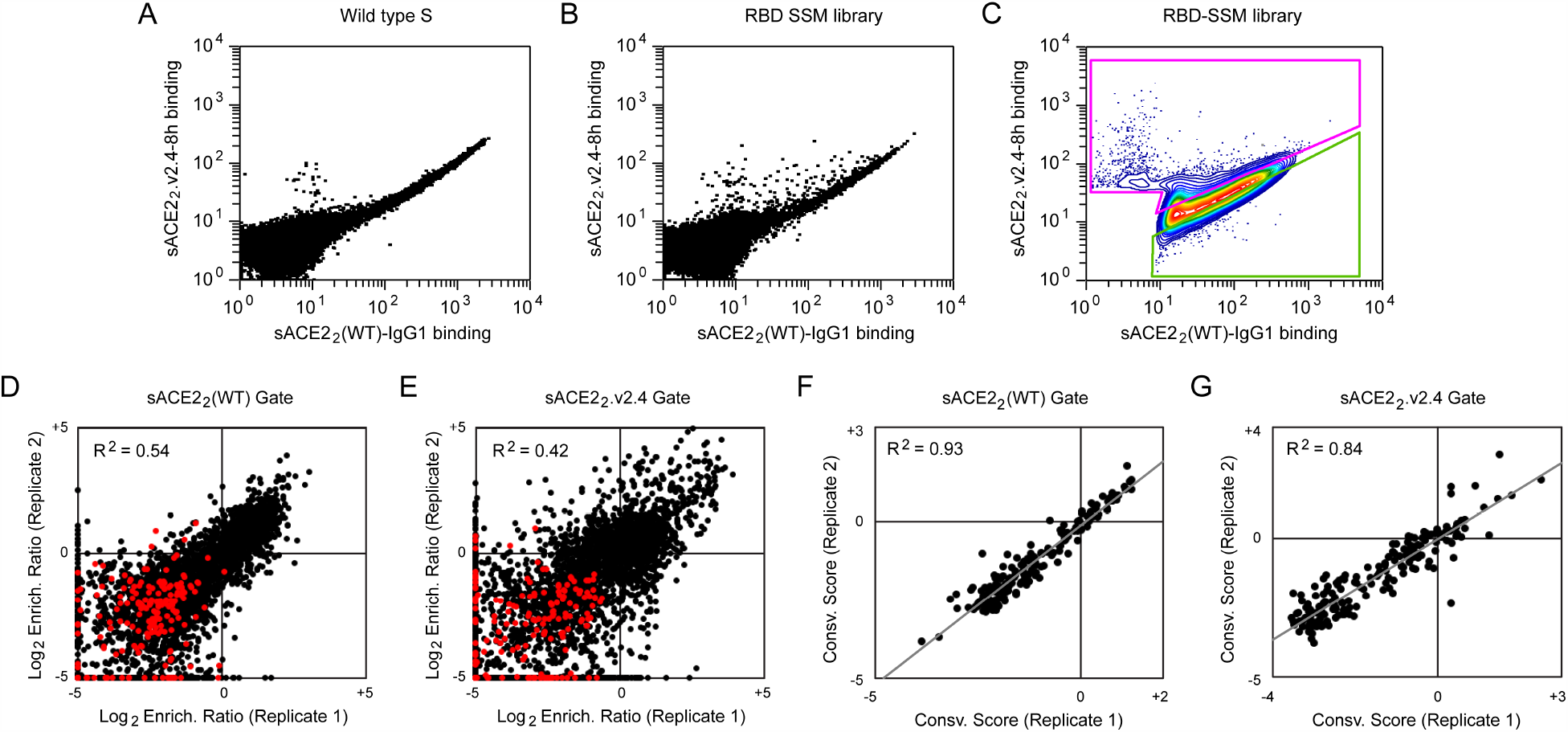
A competition-based selection to identify RBD mutations within S of SARS-CoV-2 that preferentially bind wild type or engineered ACE2 receptors. **(A)** Expi293F cells were transfected with wild type myc-S and incubated with competing sACE2_2_(WT)-IgG1 (25 nM) and sACE2_2_.v2.4-8h (20 nM). Bound protein was detected by flow cytometry after immuno-staining for the respective epitope tags. **(B)** As in A, except cells were transfected with the RBD SSM library. A population of cells expressing S variants with increased specificity towards sACE2_2_.v2.4 is apparent (cells shifted to the upper-left of the main population). **(C)** Gates used for FACS of cells expressing the RBD SSM library. After excluding cells without bound protein, the top 20% of cells for bound sACE2_2_.v2.4-8h (magenta gate) and for bound sACE2_2_(WT)-IgG1 (green gate) were collected. **(D-E)** Agreement between log_2_ enrichment ratios from two independent FACS selections for cells expressing S variants with increased specificity for (D) sACE2_2_(WT) or (E) sACE2_2_.v2.4. R^2^ values are calculated for nonsynonymous mutations (black). Nonsense mutations are red. **(F-G)** Conservation scores are calculated from the mean of the log_2_ enrichment ratios for all nonsynonymous substitutions at a given residue position. Correlation plots show agreement between conservation scores for two independent selections for cells within the (D) sACE2_2_(WT) or (E) sACE2_2_.v2.4 specific gates.

Soluble ACE2_2_.v2.4 has three mutations from wild type ACE2: T27Y buried within the RBD interface, and L79T and N330Y at the interface periphery (Figure 5A). A substantial number of mutations in the RBD of S were selectively enriched for preferential binding to sACE2_2_.v2.4 (Figure 5B, upper-left quadrant). While sACE2_2_.v2.4-specificity mutations could be found immediately adjacent to the sites of engineered mutations in ACE2 (in particular mutations to S-F486 adjacent to ACE2-L79 and S-T500 adjacent to ACE2-N330), major hot spots for sACE2_2_.v2.4-specificity mutations were also mapped to RBD loop 498-506, contacting the region where the ACE2-α1 helix packs against a β-hairpin motif (Figure 5A). By comparison, there were no hot spots in the RBD for sACE2_2_(WT)-specificity mutations. Indeed, only a small number of mutations were selectively enriched for preferential binding to wild type receptor (Figure 5B), and the abundance of these putative wild type-specific mutations barely rose above the expected level of noise in the deep mutagenesis data. In this competition assay, S binding to wild type sACE2_2_ is therefore more sensitive to RBD mutations than S binding to engineered sACE2_2_.v2.4.

**Figure 5.**
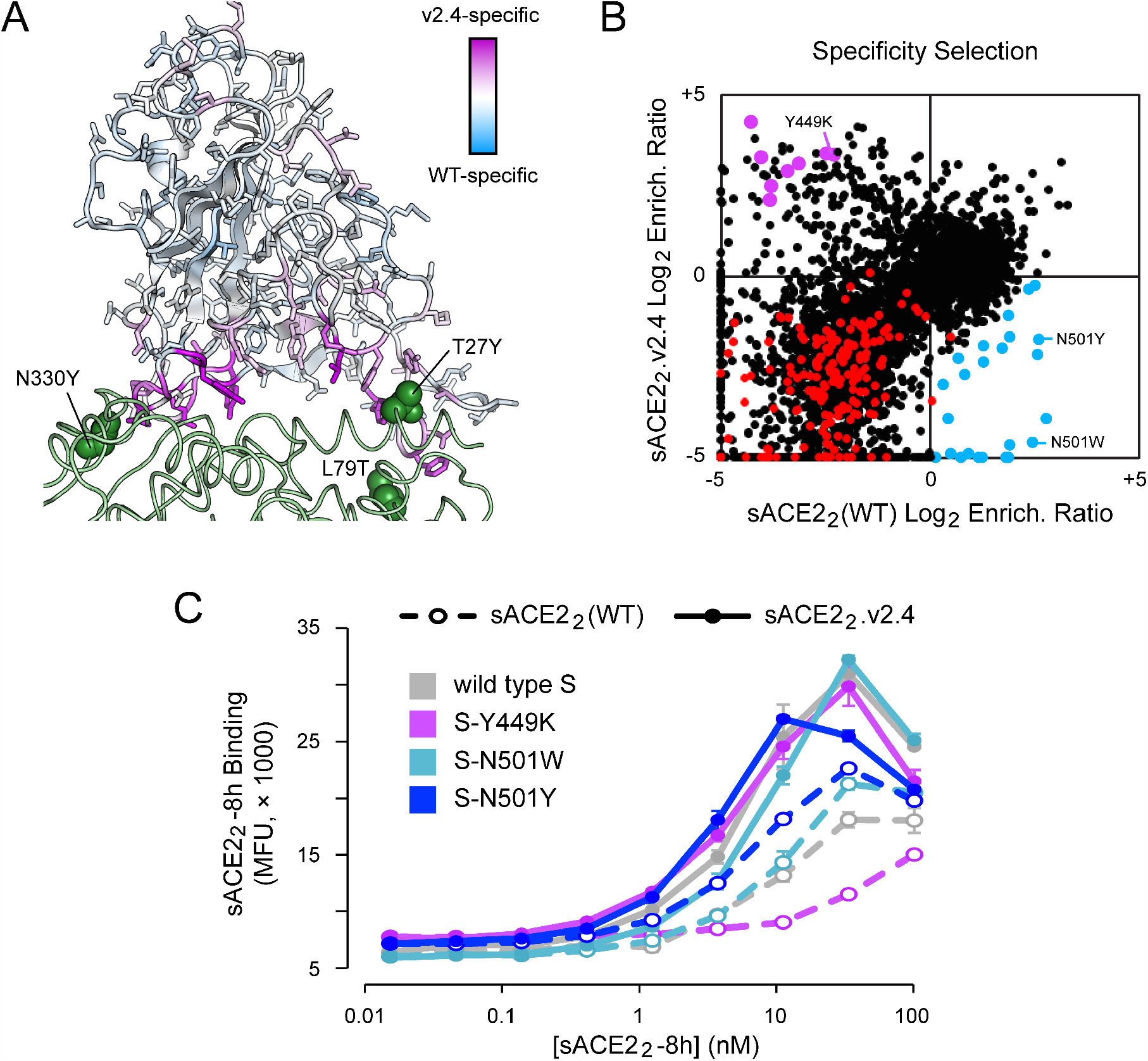
Mutations within the RBD that confer specificity towards wild type ACE2 are rare. **(A)** The SARS-CoV-2 RBD is colored by specificity score (the difference between the conservation scores for cells collected in the sACE2_2_(WT) and sACE2_2_.v2.4 specific gates). Residues that are hot spots for mutations with increased specificity towards sACE2_2_(WT) are blue or towards sACE2_2_.v2.4 are purple. The contacting surface of ACE2 is shown as a green ribbon, with sites of mutations in sACE2_2_.v2.4 labeled and shown as green spheres. **(B)** Log_2_ enrichment ratios for mutations in S expressed by cell populations collected in the sACE2_2_(WT) (x-axis) and sACE2_2_.v2.4 (y-axis) specific gates. Data are the mean from two independent sorting experiments. S mutants in blue were predicted to have increased specificity for sACE2_2_(WT) and were tested by targeted mutagenesis in Figure S6. S mutants in purple were predicted to have increased specificity for sACE2_2_.v2.4 and were tested by targeted mutagenesis in Figure S7. Other nonsynonymous mutations are black. Nonsense mutations are red. **(C)** Wild type myc-S (grey) and three variants, Y449K (purple), N501W (light blue), and N501Y (dark blue), were expressed in Expi293F cells and tested by flow cytometry for binding to sACE2_2_(WT)-8h (dashed lines) or sACE2_2_.v2.4-8h (solid lines).

To determine whether the potential wild type ACE2-specific mutations found by deep mutagenesis are real as opposed to false predictions due to data noise, we tested 24 mutants of S selectively enriched in the wild type-specific gate by targeted mutagenesis (blue data points in Figure 5B). Only minor shifts towards binding wild type sACE2_2_ were observed (Figure S6). Two S mutants were investigated further in sACE2_2_ titration experiments, N501W and N501Y, which both retained high receptor binding and displayed small shifts towards wild type sACE2_2_ in the competition experiment. N501 of S is located in the 498-506 loop and its substitution to large aromatic side chains might alter the loop conformation to cause steric strain with nearby ACE2 mutation N330Y in sACE2_2_.v2.4. After titrating the concentrations of 8his-tagged sACE2_2_(WT) and sACE2_2_.v2.4 and measuring bound protein to S-expressing cells by flow cytometry, it was found S-N501W and S-N501Y do show enhanced specificity for wild type sACE2_2_, but the effect is small and sACE2_2_.v2.4 remains the stronger binder (Figure 5C).

Dimeric sACE2_2_ binds avidly to S protein on a membrane surface; avid interactions are also observed between sACE2_2_ and spikes on authentic SARS-CoV-2 in infection assays (*27*). We used BLI kinetics measurements, in which immobilized sACE2_2_-IgG1 interacts with monomeric RBD, to determine how the observed changes in avid sACE2_2_ binding to S-expressing cells translate to changes in affinity. Both N501W and N501Y mutants of SARS-CoV-2 RBD displayed increased affinity for wild type ACE2 and engineered ACE2.v2.4, with larger affinity gains in favor of the wild type receptor (Table 1). This aligns with the flow cytometry data indicating a small shift in specificity towards wild type ACE2, but not enough to escape the engineered decoy. By comparison, multiple independent escape mutations are readily found in S of SARS-CoV-2 that diminish the efficacy of monoclonal antibodies by many orders of magnitude (*20, 21*).

Finally, 8 representative mutations to S predicted from the deep mutational scan to increase specificity towards sACE2_2_.v2.4 (purple data points in Figure 5B) were cloned and 7 were found to have large shifts towards preferential sACE2_2_.v2.4 binding in the competition assay (Figure S7). These S mutations were Y449K/Q/S, L455G/R/Y, and G504K. The basis for why the mutations increase specificity towards engineered sACE2_2_.v2.4 is ambiguous, since RBD residues Y449, L455, and G504 are not in direct contact with engineered sites of the receptor. BLI kinetics between immobilized sACE2_2_-IgG1 and monomeric RBD as the analyte showed reduced affinity of a representative mutant, RBD-Y449K, to both wild type and engineered sACE2_2_ (Table 1). However, affinity changes in the picomolar range for sACE2_2_.v2.4 are hidden during avid binding to full-length S-Y449K at the cell surface, whereas avid binding of wild type sACE2_2_ to S-Y449K (with affinity measured by BLI in the moderate nanomolar range) is substantially reduced. This finding might explain why the competition selection found many mutations that shift specificity towards engineered ACE2, as mutations causing small decreases in affinity may have larger effects on avid binding of the weaker-bound wild type receptor.

Overall, validation by targeted mutagenesis confirms that the selection can successfully find mutations in S with altered specificity. The inability to find mutations in the RBD that impart high specificity for the wild type receptor means such mutations are rare or may not even exist, at least within the receptor-binding domain where direct physical contacts with receptors occur. We cannot exclude mutations elsewhere having long-range conformational effects. Engineered, soluble decoy receptors therefore live up to their promise as broad therapeutic candidates against which a virus cannot easily escape.

## DISCUSSION

The allure of soluble decoy receptors is that the virus cannot easily mutate to escape neutralization. Mutations that reduce affinity of the soluble decoy will likely also decrease affinity for the wild type receptor on host cells, thereby coming at the cost of diminished infectivity and virulence. However, this hypothesis has not been rigorously tested, and since engineered decoy receptors differ from their wild type counterparts, even if by just a small number of mutations, it is possible a virus may evolve to discriminate between the two. Here, we show that an engineered decoy receptor for SARS-CoV-2 broadly binds with low nanomolar K_D_ to the spikes of SARS-associated betacoronaviruses that use ACE2 for entry, despite high sequence diversity within the ACE2-binding site. Mutations in S of SARS-CoV-2 that confer high specificity for wild type ACE2 were not found in a comprehensive screen of all substitutions within the RBD. The engineered decoy receptor is therefore broad against zoonotic ACE2-utilizing coronaviruses that may spill over from animal reservoirs in the future and against variants of SARS-CoV-2 that may arise as the current COVID-19 pandemic rages on. These findings are highly consistent with research of a high-affinity decoy receptor engineered for a different pathogen, the human immunodeficiency virus (HIV). An IgG1 Fc-fused soluble decoy based on HIV receptors, called eCD4-Ig, broadly neutralizes HIV-1, HIV-2, and related simian viruses, with single mutations in the HIV spike protein unable to achieve full escape (*46, 47*). Together with the results reported here, these studies collectively demonstrate that engineered decoy receptors can achieve exceptional breadth against virus sequence variants. We argue it is unlikely that decoy receptors will need to be combined in cocktail formulations, as is required for many monoclonal antibodies and possibly designed miniprotein binders to prevent the rapid emergence of resistance (*20, 48*), facilitating manufacture and distribution. Our findings give insight into how a potential therapeutic can achieve breadth with a low chance of virus resistance for a family of highly infectious and deadly viruses.

## MATERIALS AND METHODS

### Plasmids

Residue numbers for constructs begin from the start methionine as amino acid (a.a.) 1. The cloning of human codon-optimized, mature S from SARS-CoV-2 (GenBank Acc. No. YP_009724390.1; a.a. V16-T1273) into the NheI-XhoI sites of pCEP4 (Invitrogen) with an N-terminal, extracellular c-myc tag is described elsewhere (*27*). Soluble ACE2 (a.a. 1-732 encoding a dimer; wild type or engineered variant sACE2_2_.v2.4) fused to an 8his purification tag or to human IgG1-Fc (a.a. D221-K447; nG1m1 isoallotype; GenBank KY432415.1) and cloned in to the NheI-XhoI sites of pcDNA3.1(+) (Invitrogen) is also previously described (*27*). The RBDs of SARS-CoV-1 (Urbani isolate; GenBank AAP13441.1; a.a. T320-D518), SARS-CoV-2 (YP_009724390; a.a. T333-K529), LYRa11 (AHX37558.1; a.a. T324-D522), Rs7327 (ATO98218.1; a.a. T321-D519), Rs4231 (ATO98157.1; a.a. T320-D518), Rs4084 (ATO98132.1; a.a. T321-D519) and RsSHC014 (AGZ48806.1; a.a. T321-D519) were cloned with N-terminal influenza HA leader peptides (sequence MKTIIALSYIFCLVFA) and C-terminal 8-his tags (sequence GSGHHHHHHHH) into the NheI-XhoI sites of pcDNA3.1(+). These plasmids are deposited with Addgene under accession numbers 145145 and 161821-161826. Mutations were made by overlap extension PCR and verified by Sanger sequencing.

### Tissue Culture

Expi293F cells (ThermoFisher) were grown in Expi293 Expression Medium (ThermoFisher) at 125 rpm, 8 % CO2, 37 °C.

### Recombinant Protein Production

Plasmids (500 ng DNA per ml culture) and polyethylenimine (MW 25,000; Polysciences; 3 µg per ml culture) were mixed with OptiMEM (Gibco; 100 µl per ml culture), incubated 20 minutes at room temperature and added to Expi293F cells at a density of 2 × 10^6^ / ml. Transfection Enhancers (ThermoFisher) were added 18-23 h post-transfection. Culture supernatant was harvested 4-6 days later by two centrifugation steps (800 × g for 10 minutes to remove cells and 20,000 × g for 20 minutes to remove debris). IgG1 Fc fused and 8his-tagged proteins were subsequently purified as previously described (*27*) using KANEKA KanCapA 3G Affinity (Pall) and HisPur Ni-NTA (Thermo Scientific) resins, respectively. Eluted proteins from affinity chromatography were then separated on a

Superdex 200 Increase 10/300 GL column (GE Healthcare Life Sciences) equilibrated with Dulbecco’s phosphate-buffered saline (PBS). Proteins from peak fractions were concentrated using centrifugal ultrafiltration devices (Millipore) to final concentrations of ∼1 mg/ml (RBD-8h proteins), ∼10 mg/ml (sACE2_2_-8h proteins) and ∼50 mg/ml (sACE2_2_-IgG1 proteins). Concentrations were determined by absorbance at 280 nm using calculated extinction coefficients. Reported concentrations for sACE2_2_ are based on monomeric subunits. Aliquots were snap frozen in liquid N_2_ and stored at −80 °C.

### Biolayer Interferometry

BLI kinetics were collected on an Octet RED96a and analyzed with a 1:1 binding model (global fit) using instrument software (Molecular Devices). IgG1 Fc-fused sACE2_2_ (wild type or engineered variant sACE2_2_.v2.4) were immobilized at 100 nM for 10 minutes to anti-human IgG Fc biosensors (Molecular Devices). The assay buffer was 10 mM HEPES pH 7.6, 150 mM NaCl, 3 mM EDTA, 0.05% polysorbate 20, 0.5% non-fat dry milk (Bio-Rad). Loaded sensors were equilibrated for 30 s in buffer, then dipped in RBD-8h solutions for 60 s to measure association and transferred back to buffer to measure dissociation over 300 s.

### Library Construction, FACS and Illumina Sequencing Analysis

Using plasmid pCEP4-myc-S encoding tagged, full length S of SARS-CoV-2, saturation mutagenesis was focused to residues C336-L517 forming the RBD. Degenerate NNK codons were introduced at all RBD positions using overlap extension PCR as previously described (*49*). Transient transfection conditions were used that typically provide no more than a single coding variant per cell (*36, 37*). Expi293F cells at 2 × 10^6^ / ml were transfected with a mixture of 1 ng coding plasmid (i.e. library DNA) with 1.5 µg pCEP4-ΔCMV carrier plasmid (described in (*37*)). The medium was replaced 2 h post-transfection and cells were collected 24 h post-transfection for FACS. Cells were washed with ice-cold PBS supplemented with 0.2 % bovine serum albumin (PBS-BSA).

For investigations of binding to wild type sACE2_2_, the cells expressing the S library were resuspended in 2.5 nM sACE2_2_(WT)-8h and incubated 30 minutes on ice. Cells were washed twice with PBS-BSA and then co-stained for 20 minutes with anti-myc Alexa 647 (clone 9B11, 1/250 dilution; Cell Signaling Technology) and anti-HIS-FITC (chicken polyclonal, 1/100 dilution; Immunology Consultants Laboratory). Cells were again washed twice before sorting on a BD FACS Aria II at the Roy J. Carver Biotechnology Center. Dead cells, doublets and debris were excluded by first gating on the main population by forward/side scattering and then excluding DAPI positive cells. From the myc-S-positive (Alexa 647) population, the 20 % of cells with the highest and 20% of cells with the lowest anti-HIS-FITC fluorescence for bound sACE2_2_(WT)-8h were collected (Figure 2C). Collection tubes were coated overnight with fetal bovine serum prior to sorting and contained Expi293 Expression Medium. Fluorescent signals decreased over time during FACS, and therefore transfected cultures were prepared on three separate occasions for a combined total of 8 hours sort time. The total numbers of collected cells were 57,000 and 72,700 for the ACE2-High and ACE2-Low gates, respectively. Collected cells were centrifuged (500 × g, 300 s) and pellets were frozen at −80°C. Samples from the independent sorts were pooled during extraction of total RNA.

The competition selection was performed similarly, with the exception that cells expressing the S library were incubated for 30 minutes in a mixture of 20 nM sACE2_2_.v2.4-8h and 25 nM sACE2_2_(WT)-IgG1. After washing twice, bound proteins were stained for 30 minutes with anti-human IgG-APC (clone HP6017, 1/250 dilution; BioLegend) and anti-HIS-FITC (chicken polyclonal, 1/100 dilution; Immunology Consultants Laboratory). Cells were washed twice and sorted. After gating for the main population of viable cells as described above, the 20 % of cells with the highest FITC-relative-to-APC and highest APC-relative-to-FITC signals were collected (Figure 4C). The total numbers of collected cells were 53,950 (Replicate 1: sACE2_2_(WT)-specific gate), 42,860 (Replicate 1: sACE2_2_.v2.4-specific gate), 41,420 (Replicate 2: sACE2_2_(WT)-specific gate), and 34,730 (Replicate 2: sACE2_2_.v2.4-specific gate).

Total RNA was extracted from the collected cells using a GeneJET RNA purification kit (Thermo Scientific). First strand cDNA was synthesized with Accuscript (Agilent) primed with a gene-specific oligonucleotide. The region of S scanned by saturation mutagenesis was PCR amplified as 3 overlapping fragments that together span the full RBD sequence. Following a second round of PCR, primers added adapters for annealing to the Illumina flow cell and sequencing primers, together with barcodes for experiment identification. The PCR products were sequenced on an Illumina NovaSeq 6000 using a 2×250 nt paired end protocol. Data were analyzed using Enrich (*38*), where the frequencies of S variants in the transcripts of the sorted populations were compared to their frequencies in the naive plasmid library. Log_2_ enrichment ratios for all the individual mutations were calculated and normalized by subtracting the log_2_ enrichment ratio for the wild type sequence across the same PCR-amplified fragment. Conservation scores at residue positions were calculated by averaging the log_2_ enrichment ratios for all non-synonymous mutations at the residue.

### Flow Cytometry Analysis of SARS-CoV-2 S Mutants

Expi293F cells at 2.0 × 10^6^ cells/ml were transfected with plasmid DNA (300 ng per ml of culture for measuring myc-S surface expression and sACE2_2_ competition binding, 500 ng per ml for titration experiments) encoding myc-S variants using Expifectamine (ThermoFisher) according to the manufacturer’s directions. At 24 h post-transfection, cells were washed with PBS-BSA. To detect surface expressed myc-S, cells were incubated with anti-myc Alexa 647 (clone 9B11, 1/250 dilution; Cell Signaling Technology) on a rocker at 4°C for 30 minutes. To measure competitive binding of wild type and engineered receptors, cells were instead incubated with 25 nM sACE2_2_(WT)-IgG1 and 20 nM sACE2_2_.v2.4-8h for 30 minutes at 4°C, washed twice, and stained with anti-human IgG-APC (clone HP6017, 1/250 dilution; BioLegend) and anti-HIS-FITC (chicken polyclonal, 1/100 dilution; Immunology Consultants Laboratory) secondary antibodies for 20 minutes at 4°C. Finally, in titration experiments, transfected cells were incubated for 30 minutes at 4°C with 1/3 serial dilutions of sACE22(WT)-8h or sACE22.v2.4-8h, followed by two washes and a 30 minute incubation with anti-myc Alexa 647 (clone 9B11, 1/250 dilution) and anti-HIS-FITC (chicken polyclonal, 1/100 dilution). For all experiments, cells were washed twice prior to analysis on an Accuri C6 Flow Cytometer (BD Biosciences) and data were processed with FCS Express (De Novo Software). Quantification of myc-S surface expression is detailed in Figure S5.

### Reagent and Data Availability

Plasmids for RBD protein expression are deposited with Addgene (numbers 145145 and 161821-161826). Illumina sequencing data are deposited in NCBI’s Gene Expression Omnibus (GEO) under series accession number GSE159372.

## Supporting information

Enrichment ratios from deep mutagenesis of RBD.

## ACKNOWLEDGEMENTS

The Roy J. Carver Biotechnology Center at the University of Illinois assisted with flow cytometry and Illumina sequencing. E.P. designed the study and completed deep mutagenesis. K.K.C. purified proteins, tested BLI kinetics. K.K.C., K.K.N. and T.J.C.T. prepared samples for flow cytometry. This work was in part supported by NIH award R01AI129719 to E.P. The University of Illinois has filed a provisional patent for engineered decoy receptors and E.P. and K.K.C. are co-founders of Orthogonal Biologics, Inc.

## Notes

### Summary of Updates

This revision includes new BLI kinetics measurements of affinities between purified RBD of SARS-CoV-2 (including mutant N501Y) and human entry receptor ACE2.

