## Supplemental Figures S1 to S7 for "An engineered decoy receptor for SARS-CoV-2 broadly binds protein S sequence variants"

### Supporting Information

Figures S1 to S7

A

```
                340       350       360       370       380       390       400       410       420       430
                 |         |         |         |         |         |         |         |         |         |
SARS-2    TNLCPFGEVFNATRFASVYAWNRKRISNCVADYSVLYNSASFSTFKCYGVSPTKLNDLCFTNVYADSFVIRGDEVRQIAPGQTGKIADYNYKLPDDFTGC
SARS-1    TNLCPFGEVFNATKFPSVYAWERKKISNCVADYSVLYNSTFFSTFKCYGVSATKLNDLCFSNVYADSFVVKGDDVRQIAPGQTGVIADYNYKLPDDFMGC
Rs4084    TNLCPFGEVFNATTFPSVYAWERKRISNCVADYSILYNSTSFSTFKCYGVSATKLNDLCFSNVYADSFVVKGDDVRQIAPGQTGVIADYNYKLPDDFLGC
RsSHC014  TNLCPFGEVFNATTFPSVYAWERKRISNCVADYSVLYNSTSFSTFKCYGVSATKLNDLCFSNVYADSFVVKGDDVRQIAPGQTGVIADYNYKLPDDFLGC
Rs4231    TNLCPFGEVFNATTFPSVYAWERKRISNCVADYSVLYNSTSFSTFKCYGVSATKLNDLCFSNVYADSFVVKGDDVRQIAPGQTGVIADYNYKLPDDFLGC
LYRa11    TNLCPFGEVFNATTFPSVYAWERKRISNCVADYSVLYNSTSFSTFKCYGVSAIKLNDLCFSNVYADSFVVKGDDVRQIAPGQTGVIADYNYKLPDDFMGC
Rs7327    TNLCPFGEVFNATTFPSVYAWERKRISNCVADYSVLYNSTSFSTFKCYGVSATKLNDLCFSNVYADSFVVKGDDVRQIAPGQTGVIADYNYKLPDDFMGC
                                                                                *             *

                440       450       460       470       480       490       500       510       520
                 |         |         |         |         |         |         |         |         |
SARS-2    VIAWNSNNLDSKVGGNYNYLYRLFRKSNLKPFERDISTEIYQAGSTPCNGVEGFNCYFPLQSYGFQPTNGVGYQPYRVVVLSFELLHAPATVCGPK
SARS-1    VLAWNTRNIDATSTGNYNYKYRYLRHGKLRPFERDISNVPFSPDGKPCT-PPALNCYWPLNDYGFYTTTGIGYQPYRVVVLSFELLNAPATVCGPK
Rs4084    VLAWNTNSKDSSTSGNYNYLYRWVRRSKLNPYERDLSNDIYSPGGQSCS-AVGPNCYNPLRPYGFFTTAGVGHQPYRVVVLSFELLNAPATVCGPK
RsSHC014  VLAWNTNSKDSSTSGNYNYLYRWVRRSKLNPYERDLSNDIYSPGGQSCS-AVGPNCYNPLRPYGFFTTAGVGHQPYRVVVLSFELLNAPATVCGPK
Rs4231    VLAWNTNSKDSSTSGNYNYLYRWVRRSKLNPYERDLSNDIYSPGGQSCS-AIGPNCYNPLRPYGFFTTAGVGHQPYRVVVLSFELLNAPATVCGPK
LYRa11    VLAWNTRNIDATSSGNFNYKYRSLRHGKLRPFERDISNVPFSPDGKPCT-PPAFNCYWPLNDYGFYTTNGIGYQPYRVVVLSFELLNAPATVCGPK
Rs7327    VLAWNTRNIDATSTGNYNYKYRSLRHGKLRPFERDISNVPFSPDGKPCT-PPAFNCYWPLNDYGFFTTNGIGYQPYRVVVLSFELLNAPATVCGPK
                       ** *   * **                * ****      *** **  *************
```

B

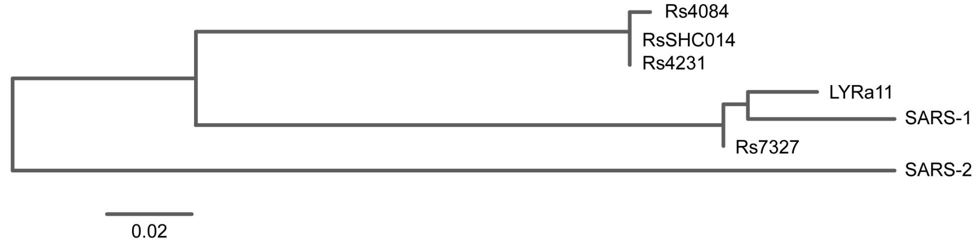


**Figure S1. The ACE2-binding site of SARS-associated betacoronaviruses is a region of high sequence diversity. (A)** RBD sequences from 2 human and 5 bat betacoronaviruses that use ACE2 as an entry receptor are aligned. Conserved residues are grey, while residues that differ are colored green (highly similar), blue (weakly similar) or red (dissimilar). Numbering is based on SARS-CoV-2 protein S. Asterisks indicate residues of SARS-CoV-2 RBD that are within 6.0 Å of ACE2 in PDB 6M17. The alignment was generated by the Clustal W server hosted by https://npsa-prabi.ibcp.fr. **(B)** Divergence (scale bar represents 2%) between RBD sequences is represented by a phylogenetic tree. Generated by the Phylogeny.fr server.

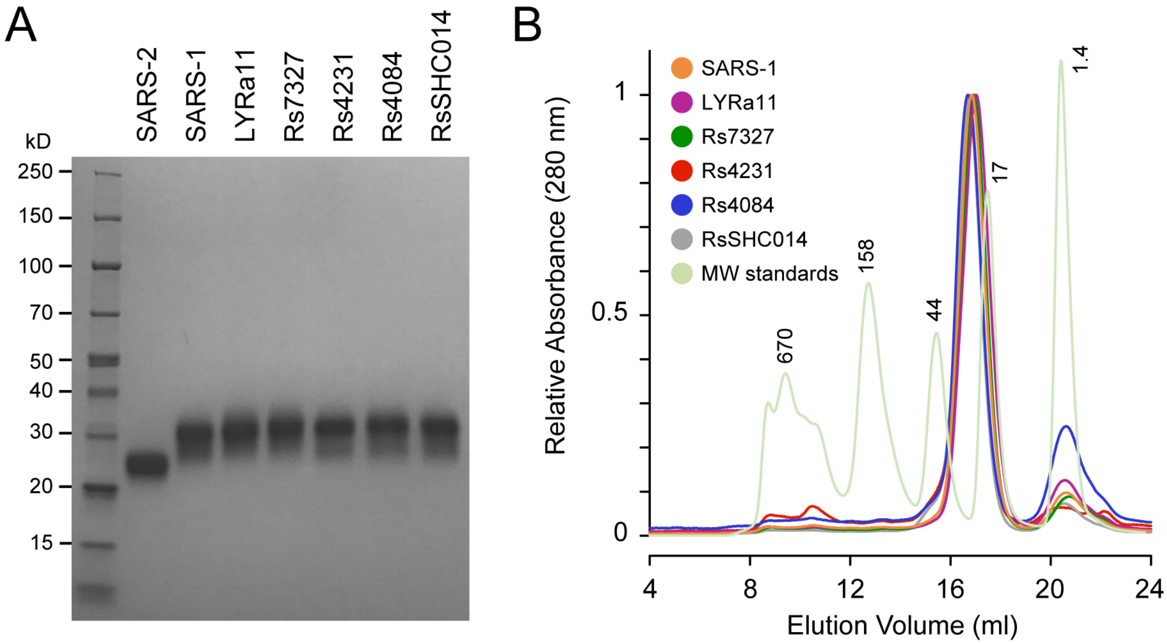


**Figure S2. Purified RBD proteins are monomers. (A)** Purified RBDs (10 μg) from the indicated viruses were separated by electrophoresis on a 4-20% SDS polyacrylamide gel and stained with Coomassie. The calculated molecular weights (MW) of RBDs, excluding any glycosylation, are ~24 kD. **(B)** Preparative size exclusion chromatography of eluted fractions from NiNTA affinity capture of 8his-tagged RBDs. UV absorbance (280 nm) is scaled and MW of standards is indicated above the peaks in kD. Peak fractions corresponding to monomeric proteins were collected for BLI kinetics analysis. The purification of SARS-CoV-2 RBD is previously reported (ref *25*).

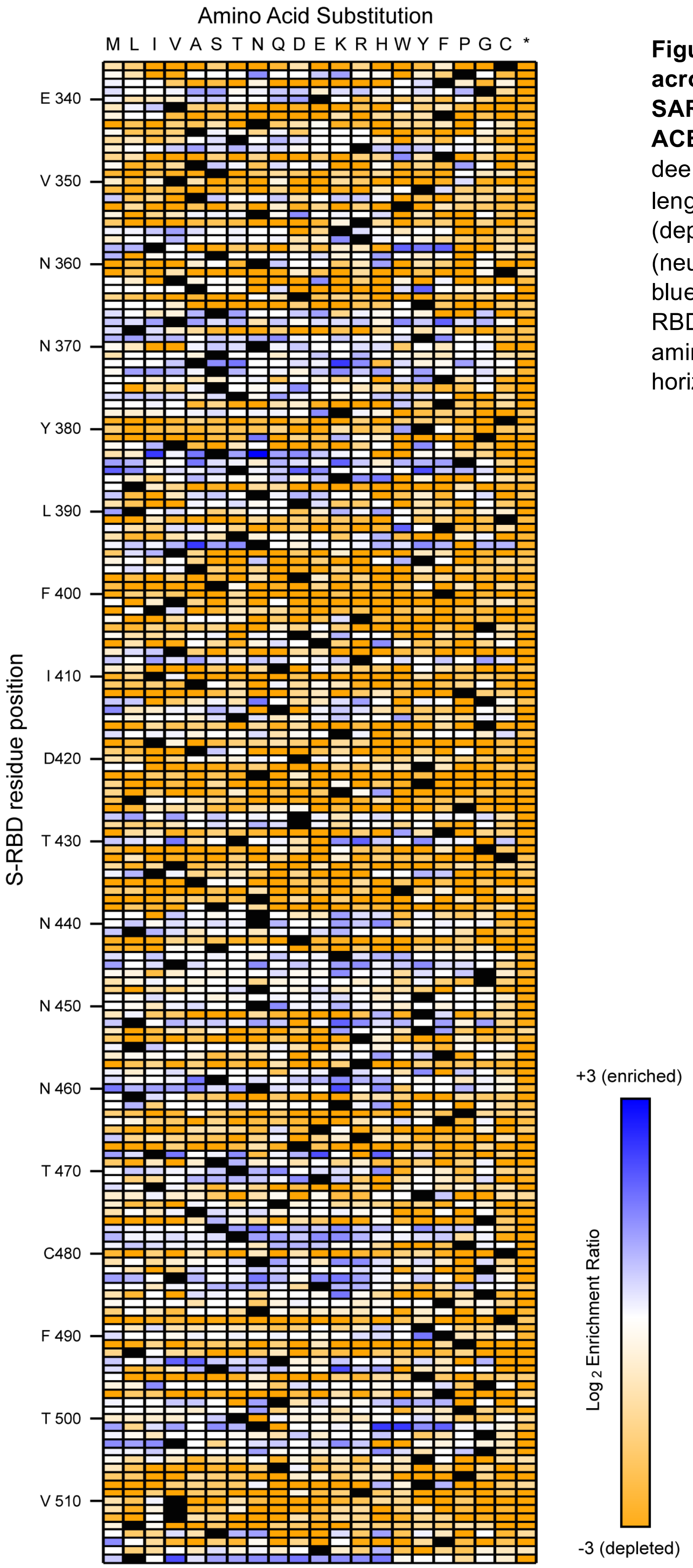


**Figure S3. The mutational landscape across the RBD of full-length S from SARS-CoV-2 for binding to soluble ACE2$_2$.** Log$_2$ enrichment ratios from the deep mutational scan of the RBD in full-length S are plotted from ≤ -3 (depleted/deleterious, orange) to 0 (neutral, white) to ≥ +3 (enriched, dark blue). Wild type amino acids are black. RBD sequence is on the vertical axis and amino acid substitutions are on the horizontal axis. *, stop codons.

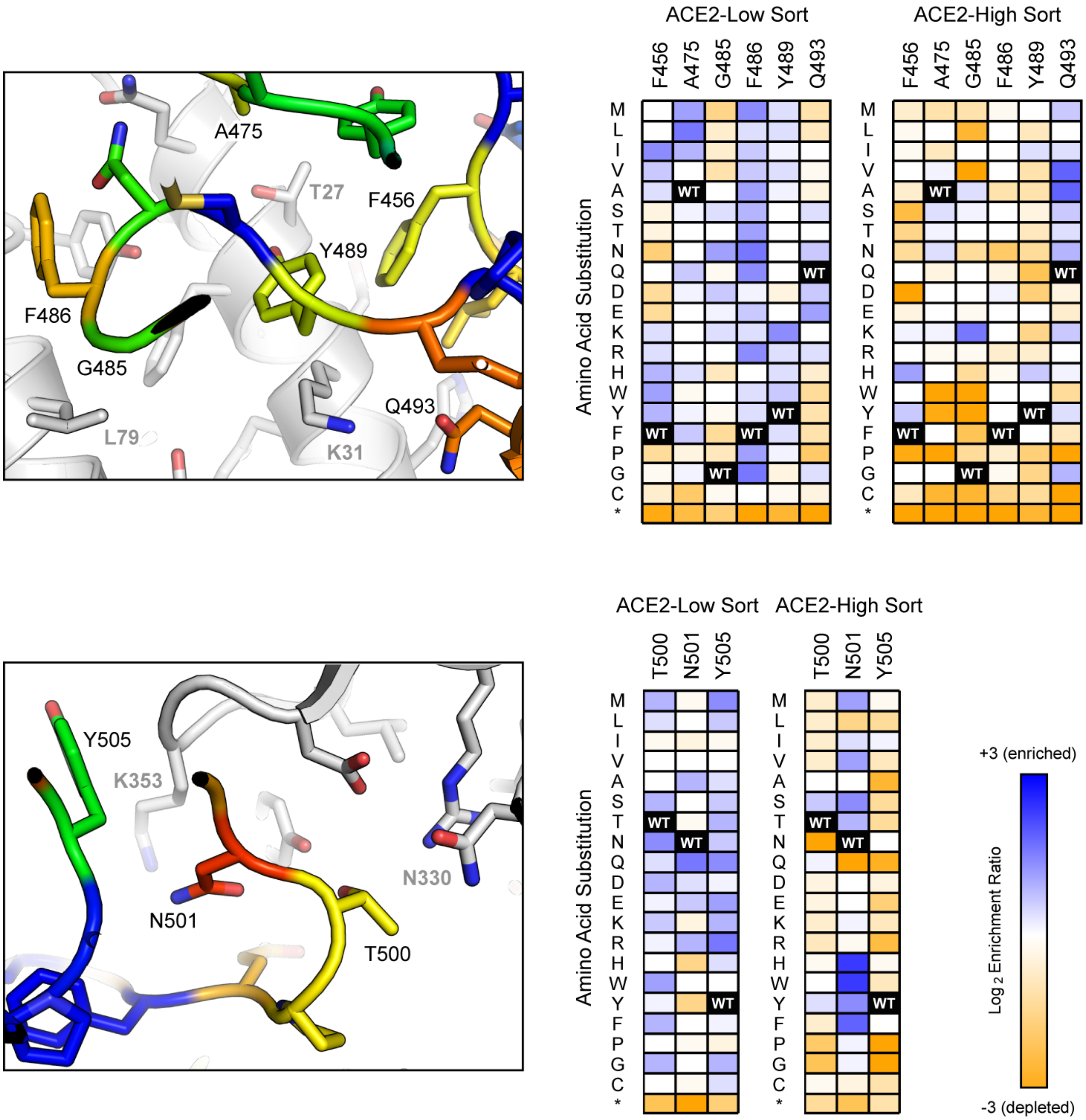


**Figure S4. SARS-CoV-2 residues at the RBD•ACE2 interface are weakly conserved for tight receptor binding.** Structural views at left (PDB 6M17) of the interface are focused on two 'hot spot' regions for interactions, centered on ACE2 residues K31 and K353. There are three substitutions in engineered sACE2.v2.4: T27Y and L79T (upper panel), and N330Y (lower panel). ACE2 residues are grey. RBD residues (labeled with black text) are colored by mutational tolerance in the ACE2-High sort experiment, from low in blue to high in red. The heatmaps at right plot the $\log_2$ enrichment ratios for S mutations, from depleted in orange to enriched in blue. Wild type amino acids are black and substitutions are indicated on the vertical axes. Mutations of SARS-CoV-2 S with increased or decreased binding to ACE2 will be preferentially enriched/blue or depleted/orange in the ACE2-High sort, respectively. Mutations of S with decreased surface expression will be depleted/orange from both the ACE2-Low and ACE2-High sorts.

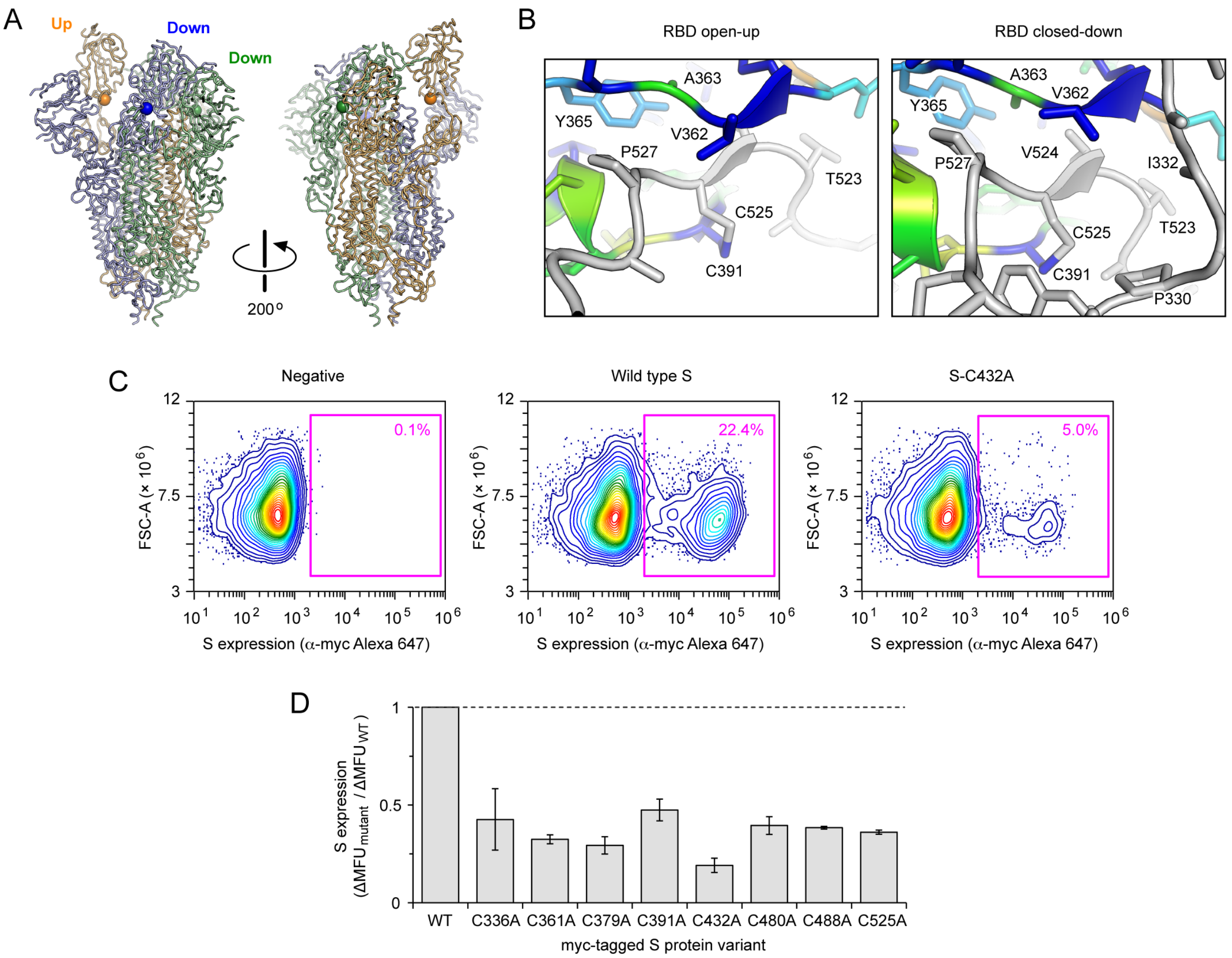


**Figure S5. Alanine substitutions of disulfide-bonded cysteines in the RBD diminish S surface expression on human cells. (A)** Structure of trimeric S (PDB 6VSB) with one subunit in a RBD open-up conformation (orange) and two subunits in closed-down conformations (blue and green). For reference, the C391 Cβ atoms are shown as spheres. **(B)** RBDs are colored by expression score from deep mutagenesis (conserved, blue; mutationally tolerant, red). Residues V362 and C391 are exposed to solvent in the open-up conformation (at left), but buried in a continuous hydrophobic core with the rest of the S1 subunit (grey) in the closed-down conformation (at right). **(C)** Based on surface immuno-staining and flow cytometry analysis, Expi293F cells transfected with myc-S cysteine mutants displayed decreases in both the percent of myc-positive cells (magenta gate) and in mean fluorescence of the positive population. To capture both effects in a single number, we calculate the change in mean fluorescence units (ΔMFU) compared to vector-transfected control cells for the entire cell population, after first gating by scattering for viable cells. **(D)** Surface expression of myc-S cysteine mutants relative to wild type myc-S. Data are mean ± range, n = 2 independent replicates.

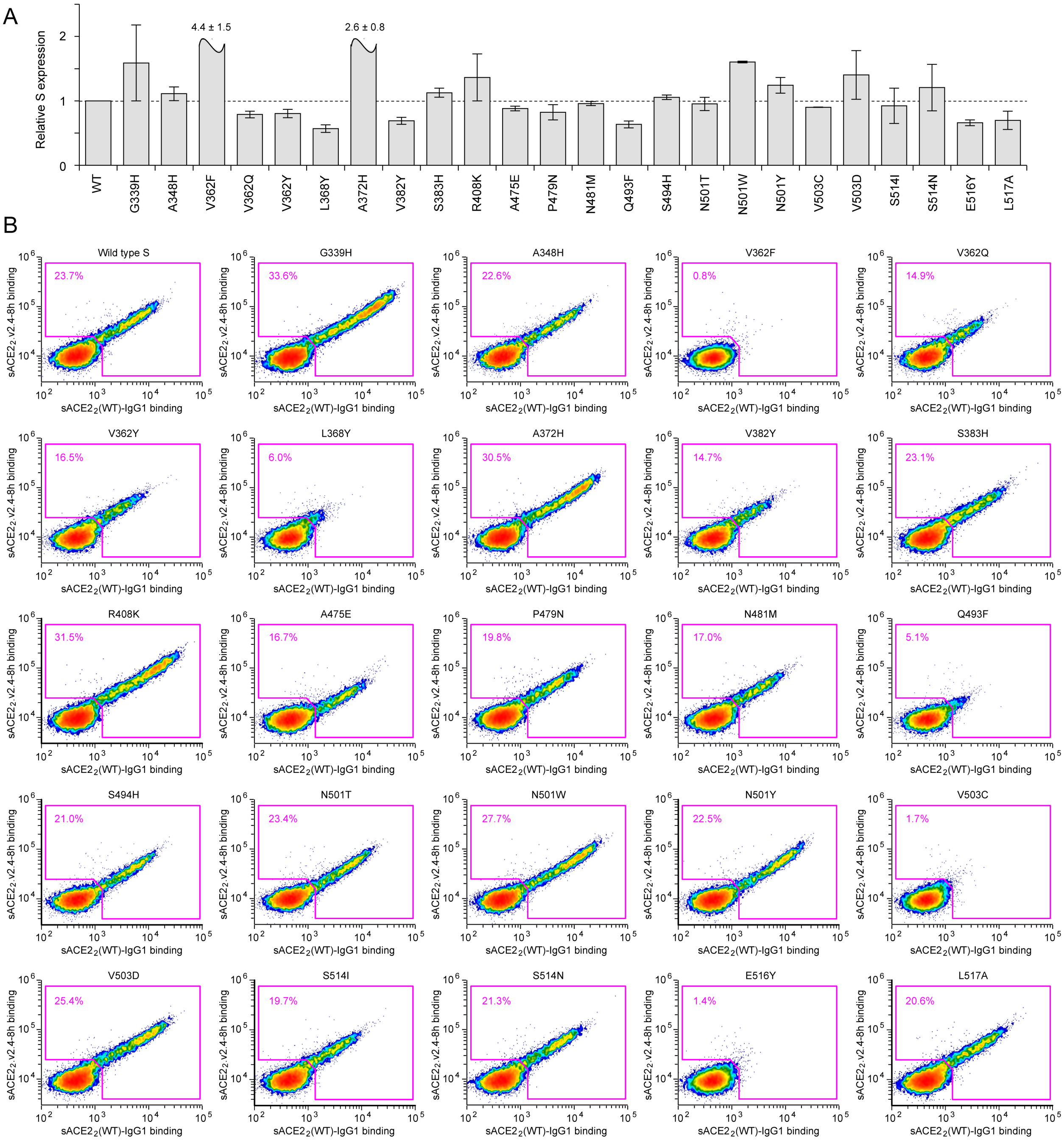


**Figure S6. Screening mutations of SARS-CoV-2 S predicted by deep mutagenesis to have enhanced specificity towards wild type sACE2$_2$ over sACE2$_2$.v2.4. (A)** Relative surface expression of myc-S mutants, determined as described in Figure S5. Data are mean ± range, n = 2 independent replicates. **(B)** Competition binding between sACE2$_2$(WT)-IgG1 (x-axis) and sACE2$_2$.v2.4-8h (y-axis) on Expi293F cells expressing the indicated myc-tagged S proteins. Cells expressing S variants with increased specificity towards wild type receptor will be shifted to the lower-right; only minor shifts are observed. Cells expressing S variants with reduced surface expression and/or ACE2 affinity have a lower percentage in the magenta gate. Results are representative of 2 replicates.

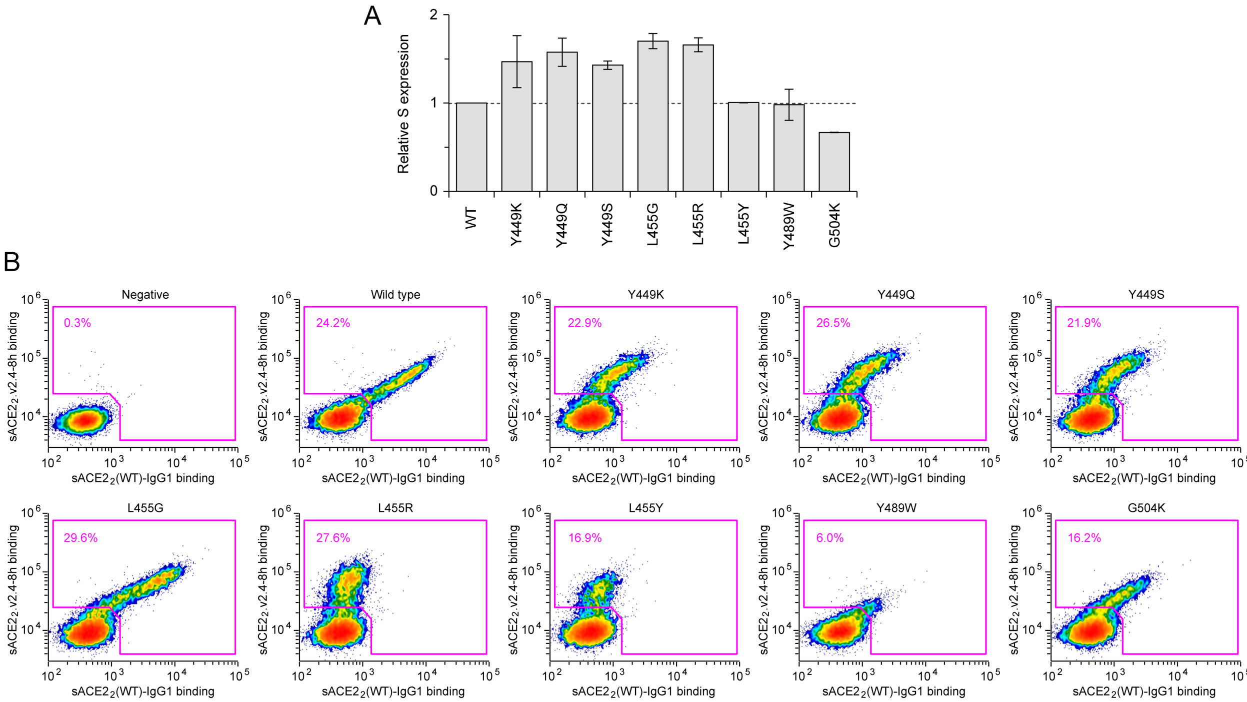


**Figure S7. Screening mutations of SARS-CoV-2 S predicted by deep mutagenesis to have enhanced specificity towards sACE2$_2$.v2.4 over wild type sACE2$_2$. (A)** Relative surface expression of myc-S mutants measured by flow cytometry. Data are mean ± range, n = 2. **(B)** Flow cytometry analysis of cells expressing myc-S variants bound to competing sACE2$_2$(WT)-IgG1 (x-axis) and sACE2$_2$.v2.4-8h (y-axis). Cells expressing S with increased specificity towards sACE2$_2$.v2.4 will be shifted to the upper-left. Results are representative of 2 replicates.
